## Supplementary material for "Inflammasomes primarily restrict cytosolic *Salmonella* replication within human macrophages": SI Figure Legends

Supplemental Figure Legends

**Figure 1—Figure Supplement 1. Caspase activity promotes the control of *Salmonella* replication within human macrophages**. WT THP-1 monocyte-derived macrophages were primed with 100 ng/mL Pam3CSK4 for 16 hours. One hour prior to infection, cells were treated with 20 µM of the pan-caspase inhibitor Z-VAD(OMe)-FMK, 25 µM of the caspase-1 inhibitor Ac-YVAD-cmk, or DMSO as a vehicle control. Then, cells were infected with PBS (Mock), WT *S*. Typhimurium (A, B, C), or WT *S.* Typhimurium constitutively expressing GFP (D) at an MOI = 20. (A) Release of IL-1β into the supernatant was measured by ELISA at 6 hpi. (B) Cell death (percentage cytotoxicity) was measured by lactate dehydrogenase release assay and normalized to mock-infected cells at 6 hpi. (C) Cells were lysed at 1 hpi and 6 hpi, and bacteria were subsequently plated to calculate CFU. Fold-change in CFU/well was calculated. (D) Cells were fixed at 6 hpi and stained for DAPI to label DNA (blue). The number of bacteria per cell at 6 hpi was scored by fluorescence microscopy. Each small dot represents one infected cell. 150 infected cells were scored for each genotype. Bars represent the mean for each condition, and error bars represent the standard deviation of triplicate wells from one experiment. ns – not significant, ***p < 0.001, ****p < 0.0001 by Dunnett's multiple comparisons test. Data shown are representative of at least three independent experiments.

**Figure 1—Figure Supplement 2. Caspase-1 promotes the control of *Salmonella* replication within human macrophages**. WT and two independent clones of *CASP1*$^{-/-}$ THP-1 monocyte-derived macrophages were primed with 100 ng/mL Pam3CSK4 for 16 hours. Cells were then infected with PBS (Mock), WT *S*. Typhimurium at an MOI = 20. (A, B, C) Release of IL-18, IL-1α, and TNF-α into the supernatant were measured by ELISA at 6 hpi. (D, E) Cells were lysed at 1 hpi and 6 hpi, and bacteria were subsequently plated to calculate CFU. (D) CFU/well at 1 hpi. (E) CFU/well at 6 hpi. Bars represent the mean for each genotype, and error bars represent the standard deviation of triplicate wells from one experiment. ns – not significant, ***$p < 0.001$, ****$p < 0.0001$ by Dunnett's multiple comparisons test. Data shown are representative of at least three independent experiments.

**Figure 1—Figure Supplement 3. Caspase-1 promotes the control of *Salmonella* replication within unprimed human macrophages**. WT and one independent clone of *CASP1*$^{-/-}$ THP-1 monocyte-derived macrophages were left unprimed for 16 hours. Cells were then infected with WT *S*. Typhimurium at an MOI = 20. Cells were lysed at 1 hpi and 6 hpi, and bacteria were subsequently plated to calculate CFU. (A) CFU/well at 1 hpi. (B) CFU/well at 6 hpi. (C) Fold-change in CFU/well was calculated. Bars represent the mean for each genotype, and error bars represent the standard deviation of triplicate wells from one experiment. ns – not significant, **$p < 0.01$ by unpaired t-test. Data shown are representative of three independent experiments.

**Figure 1—Figure Supplement 4. Stationary phase *Salmonella* does not replicate effectively in human macrophages**. WT and one independent clone of *CASP1*$^{-/-}$ THP-1 monocyte-derived macrophages were primed with 100 ng/mL Pam3CSK4 for 16 hours. Cells were then infected with stationary phase WT *Salmonella* at an MOI = 20.

Cells were lysed at 1 hpi and 6 hpi, and bacteria were subsequently plated to calculate CFU. (A) CFU/well at 1 hpi. (B) CFU/well at 6 hpi. (C) Fold-change in CFU/well was calculated. ns – not significant by unpaired t-test. Bars represent the mean for each genotype, and error bars represent the standard deviation of triplicate wells from one experiment. Data shown are representative of three independent experiments.

**Figure 1—Figure Supplement 5. Gentamicin does not contribute to the control of *Salmonella* replication in human macrophages**. WT and one independent clone of *CASP1*$^{-/-}$ THP-1 monocyte-derived macrophages were primed with 100 ng/mL Pam3CSK4 for 16 hours. Cells were then infected with WT *Salmonella* at an MOI = 20. . At 30 minutes post-infection, cells were treated with either 25 μg/ml or 100 μg/ml of gentamicin. For the 25 μg/ml gentamicin condition, at 1 hpi, cells were washed extensively with RPMI to remove the gentamicin, and then the media was replaced with fresh media containing no gentamicin. For the 100 μg/ml gentamicin condition, at 1 hpi, cells were washed extensively with RPMI to remove the gentamicin, and then the media was replaced with fresh media containing 10 μg/ml gentamicin. At 1 hpi and at 6 hpi the cells were lysed, and bacteria were subsequently plated to calculate CFU. (A) CFU/well at 1 hpi. (B) CFU/well at 6 hpi. (C) Fold-change in CFU/well was calculated. ns – not significant, ***p < 0.001 by Dunnett's multiple comparisons test. Bars represent the mean for each genotype, and error bars represent the standard deviation of triplicate wells from one experiment. Data shown are representative of three independent experiments.

**Figure 2—Figure Supplement 1. Validation of *CASP4*$^{-/-}$ THP-1 clones generated with CRISPR/Cas9-mediated genome editing**. (A) Schematic representations of the *CASP4* gene with exons (filled boxes) and introns (lines) The guide RNA target sequence for *CASP4* is highlighted in red. (B, C) Shown are the mutations of the two alleles for *CASP4* clone #2 or *CASP4* clone #6 THP-1 genomic DNA by electropherogram and sequence alignment with WT THP-1 genomic DNA. The *CASP4* target sequence is underlined. Nucleotide deletions are indicated by the red filled-in boxes. Nucleotide insertion or switch is marked by a red outline. Purple text represents the predicted impact of the mutation on the amino acid sequence. (D, E) qRT-PCR was performed to quantitate *CASP4* mRNA levels in WT THP-1s and *CASP4*$^{-/-}$ THP-1s. For the *CASP4*$^{-/-}$ THP-1s, *CASP4* mRNA levels were normalized to human *HPRT* mRNA levels and WT THP-1s. (F) Immunoblot analysis was performed on cell lysates for human CASP4 and β-actin as a loading control.

**Figure 2—Figure Supplement 2. Caspase-4 is dispensable for the control of *Salmonella* replication within human macrophages early during infection**. WT and two independent clones of *CASP4*$^{-/-}$ THP-1 monocyte-derived macrophages were primed with 100 ng/mL Pam3CSK4 for 16 hours. Cells were then infected with PBS (Mock), WT *S*. Typhimurium (A, B, C), or WT *S.* Typhimurium constitutively expressing GFP (D,E) at an MOI = 20. (A) Release of IL-1β into the supernatant was measured by ELISA at 6 hpi. (B) Cell death (percentage cytotoxicity) was measured by lactate dehydrogenase release assay and normalized to mock-infected cells at 6 hpi. (C) Cells were lysed at 1 hpi and 6 hpi, and bacteria were subsequently plated to calculate CFU.

Fold-change in CFU/well was calculated. (D, E) Cells were fixed at 6 hpi and stained for DAPI to label DNA (blue). (D) The number of bacteria per cell at 6 hpi was scored by fluorescence microscopy. Each small dot represents one infected cell. 150 infected cells were scored for each genotype. (E) Representative images are shown. Scale bar represents 10 μm. Bars represent the mean for each genotype, and error bars represent the standard deviation of triplicate wells from one experiment (A, B, C, D). ns – not significant, *$p < 0.05$ by Dunnett's multiple comparisons test (A, B, C, D). Data shown are representative of at least three independent experiments.

**Figure 2—Figure Supplement 3. Caspase-4 contributes to the control of *Salmonella* replication within human macrophages later during infection**. WT and two independent clones of *CASP4*$^{-/-}$ THP-1 monocyte-derived macrophages were primed with 100 ng/mL Pam3CSK4 for 16 hours. Cells were then infected with PBS (Mock), WT *S*. Typhimurium at an MOI = 20. (A, B, C) Release of IL-18, IL-1α, and TNF-α into the supernatant were measured by ELISA at 24 hpi. (D Cells were lysed at 1 hpi and 24 hpi, and bacteria were subsequently plated to calculate CFU. CFU/well of bacteria at 24 hpi. Bars represent the mean for each genotype, and error bars represent the standard deviation of triplicate wells from one experiment. ns – not significant, *$p < 0.05$ by Dunnett's multiple comparisons test. Data shown are representative of at least three independent experiments.

**Figure 3—Figure Supplement 1. GSDMD-mediated pore formation promotes the control of *Salmonella* replication within human macrophages.** (A, B,C) WT THP-1 monocyte-derived macrophages were primed with 100 ng/mL Pam3CSK4 for 16 hours. (D, E, F) Primary human monocyte-derived macrophages (hMDMs) were primed 500 ng/mL LPS for 3 hours. One hour prior to infection, cells were treated with 40 μM of disulfiram, to inhibit GSDMD pore formation, or DMSO as a vehicle control. Then, cells were infected with PBS (Mock), WT *S*. Typhimurium at an MOI = 20. (A, D) Release of IL-1β into the supernatant was measured by ELISA at 6 hpi. (B, E) Cell death (percentage cytotoxicity) was measured by lactate dehydrogenase release assay and normalized to mock-infected cells at 6 hpi. (C, F) Cells were lysed at 1 hpi and 6 hpi, and bacteria were subsequently plated to calculate CFU. Fold-change in CFU/well was calculated. (A, B, C) Bars represent the mean for each genotype, and error bars represent the standard deviation of triplicate wells from one experiment. Data shown are representative of at least three independent experiments. (D, E, F) The pooled results of three independent experiments using hMDMs from three different healthy human donors are shown. Each data point represents the mean of triplicate infected wells from an individual donor. ns – not significant, *p<0.05, **p< 0.01, ***p < 0.001, ****p < 0.0001 by unpaired t-test (A, B, C) or by paired t-test (D, E, F).

**Figure 3—Figure Supplement 2. GSDMD promotes the control of *Salmonella* replication within human macrophages.** WT and *GSDMD*$^{-/-}$ THP-1 monocyte-derived macrophages were primed with 100 ng/mL Pam3CSK4 for 16 hours. Cells were then infected with PBS (Mock), WT *S*. Typhimurium at an MOI = 20. (A, B, C) Release of IL-18, IL-1α, and TNF-α into the supernatant were measured by ELISA at 6 hpi. (D, E) Cells were lysed at 1 hpi and 6 hpi, and bacteria were subsequently plated to calculate

CFU. (D) CFU/well at 1 hpi. (E) CFU/well at 6 hpi. Bars represent the mean for each genotype, and error bars represent the standard deviation of triplicate wells from one experiment. ns – not significant, **p< 0.01, ****p < 0.0001 by Šídák's multiple comparisons test (A, B, C) or by unpaired t-test (D, E). Data shown are representative of at least three independent experiments.

**Figure 4—Figure Supplement 1. Cytoprotection by glycine promotes *Salmonella* replication within human macrophages.** WT and *NAIP*$^{-/-}$ THP-1 monocyte-derived macrophages were primed with 100 ng/mL Pam3CSK4 for 16 hours. 30 minutes prior to infection, cells were treated with 20 mM glycine, to prevent cell lysis, or distilled water, as a vehicle control. Cells were then infected with PBS (Mock) or WT *S*. Typhimurium at an MOI = 20. (A) Cell death (percentage cytotoxicity) was measured by lactate dehydrogenase release assay and normalized to mock-infected cells at 6 hpi. (B, C) Release of IL-1β and TNF-α into the supernatant was measured by ELISA at 6 hpi. (D, E, F) Cells were lysed at 1 hpi and 6 hpi, and bacteria were subsequently plated to calculate CFU. (D) CFU/well at 1 hpi. (E) CFU/well at 6 hpi. (F) Fold-change in CFU/well was calculated. Bars represent the mean for each condition. Error bars represent the standard deviation of triplicate wells from one experiment. *p < 0.05, **p< 0.01, ****p < 0.0001 by Tukey's multiple comparisons test. Data shown are representative of at least three independent experiments.

**Figure 5—Figure Supplement 1. Inflammasome activation primarily controls the cytosolic population of *Salmonella* in primary human macrophages.** (A, B) One hour prior to infection, human monocyte-derived macrophages (hMDMs) were treated with 20 μM of the pan-caspase inhibitor Z-VAD(OMe)-FMK or DMSO as a vehicle control. Then, cells were infected with PBS (Mock) or WT *S.* Typhimurium constitutively expressing mCherry and harboring the GFP cytosolic reporter plasmid, pNF101, at an MOI = 20. Cells were fixed at 8 hpi and stained for DAPI to label DNA (blue). (A) The number of GFP-positive bacteria (cytosolic) per cell and the number of GFP-negative, mCherry-positive bacteria (vacuolar) per cell were scored by fluorescence microscopy. Each small dot represents one infected cell. 100 total infected cells were scored for each genotype. (B) Representative images from 8 hpi are shown. Scale bar represents 10 μm. White arrows indicate cytosolic bacteria (GFP-positive). ns – not significant, ****$p < 0.0001$ by Tukey's multiple comparisons test (B). Bars represent the mean for each condition, and error bars represent the standard deviation of triplicate wells from one experiment (B). Data shown are representative of two independent experiments using hMDMs from two different healthy human donors (A, B).

**Figure 6—Figure Supplement 1. Characterization of *Salmonella* subpopulations in human macrophages.** (A, B) WT and *CASP1*$^{-/-}$ THP-1 monocyte-derived macrophages were primed with 100 ng/mL Pam3CSK4 for 16 hours. Cells were then infected with WT *S*. Typhimurium at an MOI = 20. At 8 hpi, cells were fixed and collected to be processed for transmission electron microscopy. (A) The percentage of cytosol-exposed *Salmonella* from the total number of bacteria was quantified in 20 infected cells per genotype. (B) Unannotated representative transmission electron micrographs are shown. (C) *CASP1*$^{-/-}$ THP-1 monocyte-derived macrophages were primed with 100 ng/mL Pam3CSK4 for 16 hours. Cells were then infected with WT *S*. Typhimurium at an MOI = 20. At 8 hpi, cells were fixed and collected to be processed for transmission electron microscopy. Unannotated representative tomogram slices are shown.
